# A test of the evolution of increased competitive ability in two invaded regions

**DOI:** 10.1101/589143

**Authors:** Michael C. Rotter, Mario Vallejo-Marin, Liza M. Holeski

## Abstract

1. Finding patterns that predict and explain the success of non-native species has been an important focus in invasion ecology. The evolution of increased competitive ability (EICA) hypothesis has been a frequently used framework to understand invasion success. Evolution of increased competitive ability predicts that 1. Non-native populations will escape from coevolved specialist herbivores and this release from specialist herbivores should result in relaxed selection pressure on specialist-related defense traits, 2. There will be a trade-off between allocation of resources for resistance against specialist herbivores and allocation to traits related to competitive ability and 3. This shift will allow more allocation to competitive ability traits.
2. We tested the predictions of EICA in the model plant *Mimulus guttatus*, a native of western North America (WNA). We compared how well the predictions of EICA fit patterns in two non-native regions, the United Kingdom (UK) and eastern North America (ENA). Coupled with extensive herbivore surveys we quantified genetic variation for herbivore resistance traits and fitness/ competitive ability traits to test adherence to the predictions of EICA in a common greenhouse environment.
3. Herbivore communities differed significantly between WNA, UK, and ENA populations with evidence of specialist herbivore escape in the UK, but not necessarily the ENA plants. Compared to native plants, resistance traits were lower in non-native UK plants with the exception of trichome density, while the non-native ENA plants had equivalent or higher levels of herbivore resistance traits. The UK plants had increased competitive traits than native plants while the ENA plants had equivalent competitive traits to native plants. The UK plants, but not the ENA plants, showed some signs of tradeoffs between resistance traits and fitness/ competitive ability.
4. *Synthesis*. Plants from the UK conformed to predictions of EICA more closely than those from ENA. The UK invasion is an older, more successful invasion, suggesting that support for EICA may be highest in more successful invasions. The lack of comprehensive conformity of either non-native region to the predictions of EICA also leaves room for other hypotheses that may add to our mechanistic understanding of the success of non-native plant invasions.

## Introduction

The translocation of non-native species into areas outside of their native range provides unique opportunities for the study of evolution (Cox, 2004), including how selection pressures from herbivores can shape plant defense evolution (Callaway & Maron, 2006). Comparisons between divergent biotic and abiotic factors in the native and non-native habitats can aid understanding of how these variables shape evolution in non-native plants (Whitney & Gabler, 2008). The testing of theories blending ecological and evolutionary explanations can provide important insight into how non-native plants are successful and how defense traits evolve; these tests often involve comparison of genotypes from the native and non-native ranges (Orians & Ward, 2010). Better understanding of the mechanisms of non-native plant success may allow improved control and/or more accurate predictions of the impacts that non-native species can have on native ecosystems.

Many hypotheses have been proposed to find a common reason for why plants become invasive (Catford et al., 2009), dating back to Darwin’s naturalization hypothesis (Diez et al., 2008). Hypotheses have speculated on the potential for non-native plants to more efficiently use resources than native plants (Coley et al., 1985), or have proposed that non-native plants are able to exploit an empty or less-crowded niche in the invaded habitat (Mack et al., 2000; Hierro et al., 2005,). Many of these hypothesis also incorporate the idea that a competitive advantage is gained through an enemy release in a non-native habitat from co-evolved, specialist herbivores present in the native species range (Keane & Crawley, 2002; Orians & Ward, 2009). According to one prominent hypothesis, the evolution of increased competitive ability (EICA) hypothesis, enemy release results in the allocation of resources to reproductive fitness and/or competitive ability traits rather than to defenses. Relaxed selection for defense traits would allow for the evolution of traits that allow plants to become more competitive and contribute to their invasive success (Blossey & Notzold, 1995). For instance, Blossey and Notzold (1995) found that plants in non-native populations of *Lythrum salicaria* (Lythraceae) in eastern North America produced more seeds and had greater biomass than those in native, European populations. Increases in seed production and biomass were correlated with a decline in defenses against two specialist herbivores that, at the time, were not present in eastern North America. By generating testable predictions of the role that ecology plays in shaping the evolution of non-native plants, EICA hypothesis has become one of the most widespread frameworks to explore the ability of non-native plants to succeed (Bossdorf et al., 2005; Orians & Ward, 2010).

Two specific, testable predictions of EICA to explain the success of non-native plant populations include: Firstly, non-native populations will escape from coevolved specialist herbivores that were present within the native range. This release from specialist herbivores should result in relaxed selection pressure on specialist-related defense traits (e.g., Vila et al., 2005). Secondly, EICA predicts that there will be a trade-off between allocation of resources for resistance against specialist herbivores and allocation to traits related to competitive ability. This shift to allow more allocation to competitive ability traits (e.g. increased reproduction, large plants, etc.) allows the non-native populations to compete successfully in their new habitat (Blossey & Notzold, 1995, Rotter & Holeski, 2018).

Experimental tests of EICA can be complicated by a number of factors, including the difficulty in knowing the most relevant defense traits against specialists, and inferring which competitive ability traits are most important in a particular non-native environment. Further, to test evolutionary trade-offs, traits must be studied in a common garden environment, as the measurement of phenotypes in the field yields trait values influenced by both genetic and environmental variation. Perhaps in part because of these complications, there has been mixed support for EICA (Bossdorf et al., 2005; Felker-Quinn et al., 2013; Rotter & Holeski, 2018). For example, in a recent meta-analysis that found some support for EICA, support was strongest when looking at actual herbivory (e.g. field damage or feeding trials), while there was very little support when studies looked directly at resistance traits (Rotter & Holeski, 2018).

While a number of studies have tested independent premises of EICA, fewer have conducted simultaneous assessment of both resistance and competitive ability-related traits in a common garden setting, which is necessary to detect evolutionary trade-offs (Rotter & Holeski, 2018). Here we test the predictions of EICA in *Mimulus guttatus*, using populations in the native range of western North America, as well as non-native populations in two areas of introduction, eastern North America and the United Kingdom. Specifically we tested for:

1. An escape in non-native populations from co-evolved specialist herbivore species present in the native western North American range. This would be supported by the lack of specialist herbivores feeding on *M. guttatus* in the non-native populations in eastern North America and/or the United Kingdom.
2. A decrease in herbivore resistance traits within the non-native populations, relative to native. This would be demonstrated by reduced levels of genetic-based herbivore resistance traits, or in increased performance of herbivores feeding on non-native, vs. native plants.
3. An increase in competitive ability within the non-native populations, relative to native. This would be demonstrated by increased trait values for fitness/ competitive ability traits in the eastern North America and/or UK populations, relative to native.
4. Genetic-based tradeoffs between herbivore resistance traits and competitive ability traits. Evidence of this would include negative correlations between resistance and competitive ability traits in non-native plants.
5. These predictions should be most closely followed by plants from a seemingly successful invasion (the United Kingdom plants that have filled available niches) than those from the less successful eastern North American invasion, which consists of relatively few small populations that have not expanded or have been locally extirpated.

## Methods

### Study system

*Mimulus guttatus* Fisch. ex DC. (*Erythranthe guttata* G.L. Nesom) is a species complex native to moist habitats throughout western North America (WNA). In the past few decades *Mimulus spp*. and in particular *M. guttatus*, have become important model organisms for the study of evolutionary ecology and genetics (Wu et al., 2008; Yuan, 2018). *Mimulus guttatus* has been introduced throughout the globe where it has escaped numerous times from cultivation. Non-native *M. guttatus* populations are located in the United Kingdom (UK), western Europe, New Zealand, and eastern North America (ENA) (Hall & Willis 2006, Vallejo-Marin & Lye, 2013). Historical records suggest that the first *M. guttatus* introduced in the United Kingdom originated from Alaska (Puzey & Vallejo-Marin, 2014). The first records of naturalized *M. guttatus* in the UK are from the first half of the 1800s and this taxon is currently widespread and locally abundant in the UK (Preston et al., 2002; Vallejo-Marin& Lye, 2013; Puzey & Vallejo-Marin, 2014). In contrast, it is unknown when *M. guttatus* was first introduced into ENA, but we found no collections before the early 1900’s and most extant populations were observed since the 1960’s. The source of the ENA populations is currently uncertain, but they likely represent a mix of multiple accidental introductions (e.g. through introduction of debris on military or construction equipment) and/or cultivated escapes (Gleason & Cronquist 1991).

The degree of invasiveness differs between the UK populations and the ENA populations. For instance, in particular areas of Europe there is concern over its spread into natural areas (Truscott et al., 2006) and new locations (Tokarska-Guzikand & Dajdok 2010). Within the United Kingdom, the presence of *M. guttatus* is associated with local declines in native species richness (Truscott et al., 2008). In contrast, many of the reported populations in ENA appear to not be spreading or have disappeared entirely (Timothy Block, personal communication, Gleason & Cronquist 1991).

### Plant material

We collected seed from wild populations in the summer of 2015, 2016, and 2017 in both the native (WNA) and the non-native (ENA and the UK) regions (Figure 1, Table S1). Populations were chosen to maximize geographic spread in all regions as well as to capture life history variation across the *M. guttatus* range (e.g., annual and perennial populations). We also grouped native populations into geographic clades (sub-regions in this paper) based on the genetic population structure results from Twyford and Friedman (2015) who found 5 broad genetic clusters that were geographically separated. In each population, we collected seeds from >20 plants separated by at least one meter to avoid clones and collected from multiple flowers on each plant. Populations were then grown in the greenhouse for at least one generation, and multiple maternal sib families (each family likely a mixture of full and half-siblings) were used from each population.

**Figure 1.**
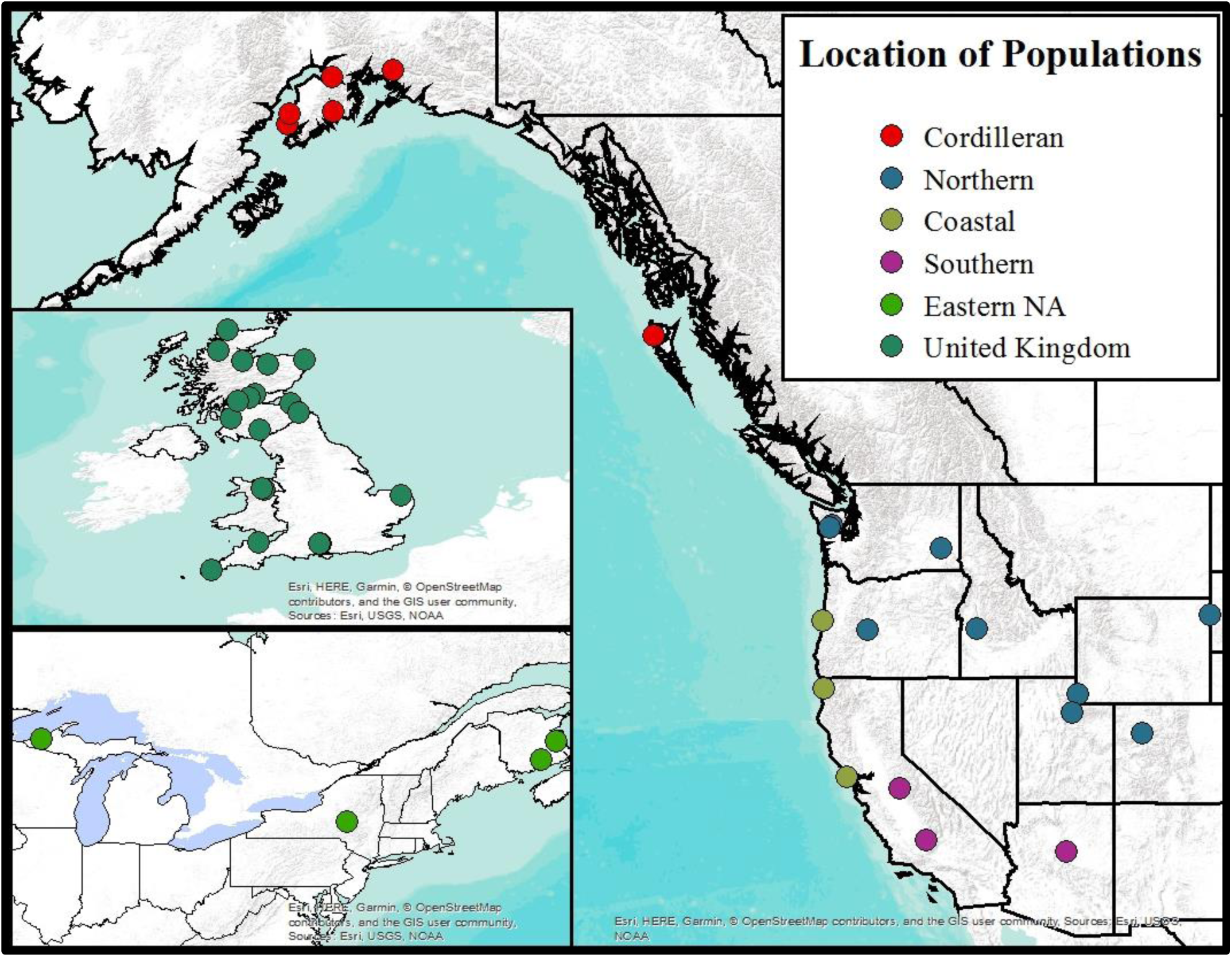
Locations of populations used in this study. Sub-regions within the native range (W. North America) are based on molecular evidence from Twyford and Friedman (2015).

### Herbivore communities

The first prediction of EICA is that there is a release from specialist herbivore pressure in non-native populations. To test for this prediction in *M. guttatus*, we collected herbivores at each seed collection site and made herbivore collections from additional populations in each region. Most sites were surveyed over at least two seasons. Plant damage was estimated at each field site as the proportion of plants in the population with visible damage measured in a discrete scale (none: no damage on any plant; low: 1-10% of plants damaged by herbivores; moderate: 10-60% of plants damaged; high: 60-90% of plants damaged; extreme: >90% of plants damaged). We also noted if the damage was caused by a mammal and any signs of what mammal species may have been responsible. For invertebrate herbivores, surveys consisted of timed visual searches and timed sweep netting (the latter only when *M. guttatus* density was high enough to preclude herbivores on other plant species). All invertebrates were collected and identified to the lowest taxonomic level possible. Herbivores were considered as those animals seen actively feeding on a plant, or those on the plant and likely able to feed on *M. guttatus* (such as a hemipteran resting on a plant but not actively feeding). In addition to these field-based surveys in both ranges, we conducted a literature review on reports of herbivores and noted their geographic range. We also looked at feeding records in the literature for herbivores of plant species closely related to *M. guttatus* (i.e., Scrophulariaceae *sensu lato*) to see if there were any specialist herbivores that may be able to shift hosts onto *M. guttatus* in the non-native regions.

### Resistance traits

Following a release from specialist herbivores, EICA predicts the evolution of lower levels of some herbivore resistance traits. To test this part of EICA we used plants derived from native and non-native populations to assess patterns of genetic- based trait variation. We assessed specific leaf area (SLA), leaf water content, leaf dry matter content (LDMC), trichome density, and foliar phytochemistry. After growing the plants in a common greenhouse environment for one month, we harvested one leaf from the fourth true-leaf pair. We weighed the leaf to get wet mass and then scanned the leaf (Epson Perfection V19) to find leaf area using Image J (Rueden et al., 2017). Freeze dried leaves (see below) were used to estimate dry weight and calculate specific leaf area (SLA), leaf water content, and leaf dry matter content (LDMC). Leaf water content and LDMC are associated with performance of some generalist herbivores consuming native *M. guttatus* (Rotter unpublished data). Trichome density was measured by counting all the trichomes at the basal section of the adaxial side of each leaf within the field of view of a dissecting microscope at 25x magnification. This density was converted to trichome density per cm^2^ (Holeski, 2007).

For phytochemical analysis, we quantified phenylpropanoid glycosides (PPGs), the predominant foliar bioactive secondary compound in the species (Holeski et al., 2013; Keefover-Ring et al., 2014). The leaf opposite the leaf in feeding trials (detailed below) was cut at the base of the petiole with scissors and flash frozen in liquid nitrogen before being transferred to a −20 degree C freezer. Tissue was then lyophilized using a pre-chilled FreeZone triad freeze dry system (Labconco; Kansas City, USA). We finely ground the freeze-dried tissue in a small capacity ball mill (dental amalgamator with steel bearings). Samples were stored and extracted as described in Holeski et al. 2013. We quantified the PPG content of each sample via high-performance liquid chromatography [HPLC; Agilent 1260 HPLC with a diode array detector and Poroshell 120 EC-C18 analytical column (4.6 · 250 mm, 2.7 μm particle size); Agilent Technologies, Santa Clara, CA] maintained at 30°C, as described in Kooyers et al. (2017). The seven PPGs analyzed in this study represent the PPGs present in detectable levels in the populations used in this study.

### Herbivore feeding trials

Herbivore response to plant resistance traits are often diffuse and vary depending on many different factors. In addition to quantifying resistance traits, we thus also measured resistance though two performance trials. For these trials, we used a subset of plant populations that represent the range of native and non-native populations (Table S1). We conducted no-choice performance trials with neonate Lepidopteran larvae of the specialist herbivore *Junonia coenia* and the generalist herbivore *Trichoplusia ni* (Rotter & Holeski, 2017; Rotter et al., 2018). One leaf from a leaf pair was placed in an envelope and treated as described above for analysis of PPGs. We assessed trichome density on the second leaf of the leaf pair, as described in Holeski (2007). The leaf scored for trichomes was then placed into a water pic and placed in a plastic container. In each container, we placed a single recently emerged first instar caterpillar. Leaves were immediately replaced with leaves from the same plant (with the opposite leaf harvested for phytochemical analysis) if/when the caterpillar consumed the entire leaf or if the leaf wilted. After larvae had fed for 10 days, we ended each trial, froze the caterpillars, and then dried and weighed them to determine caterpillar final dry mass. Larval initial (wet) weights were all within 0.001μg of each other for a particular species, so we assumed that initial dry mass was identical across larvae within each species. Higher caterpillar mass and growth rates are important indicators of greater pupal survival rates as well as increased adult fitness (Haukioja & Neuvonen, 1985; Awmack & Leather, 2002). Additionally, a more rapid growth rate allows greater survival when faced with pressure from predators and parasitoids (Feeny, 1976; Benrey & Denno, 1997).

### Plant fitness traits

Finally, EICA predicts an increase in fitness/competitive ability traits with a release from specialist herbivores and the decline of herbivore resistance traits. To test plant fitness traits we used the plants from the resistance traits measurements. We grew all plants for a total of six months prior to harvest with the exception of several populations of annual plants that were harvested after they stopped producing flowers. We assessed traits related to reproductive development, reproductive fitness, and vegetative fitness. We assessed reproductive development by counting the number of days until a plant first flowered. We also measured the corolla width (bigger flowers have been associated with pollinator preference; Martin, 2004) of the first flower on the day after it was fully emerged. We collected pollen from the first two flowers. Pollen was then stained, counted, and evaluated for viability with a hemocytometer following the procedure in Kearns and Inouye (1993). We self-pollinated each plant with the next three flowers, saturating each stigma with as much pollen as possible. Seeds were collected from these flowers and total seed was counted. Finally, the total number of flowers produced by a plant were counted at the time of plant harvest. Plants that did not flower by the end of the six-month trial (n=32 plants) were excluded from these analyses. Vegetative traits quantified included specific leaf area and leaf water content, which were measured as described above during our quantification of resistance traits. At harvest, we measured the total height (length) of the plant, from the root crown to the end of the largest shoot. We then dried all plants in a drying oven and measured aboveground biomass, belowground biomass, and total (aboveground + belowground) biomass.

### Statistical analysis

To compare herbivore communities, we used non-parametric multidimensional scaling (NMDS) to look at herbivore family and functional feeding guild differences between native and non-native populations of *M. guttatus*. The NMDS was performed using PC ORD v. 6 (McCune & Mefford, 2016). We used Jaccard distance as the similarity measure, and the program was run on “Autopilot” mode under the “slow and thorough” method, with principal axes rotation. Significance of the ordination was based on a Monte Carlo test with 250 iterations. In addition to the NMDS we looked for differences between the non-native populations and the native sub-regions in the above herbivore communities using multi-response permutation procedures (MRPP). We also used ANOVA (transformed with either a log or root transformation as assessed by Q-Q plots; we used Kruskal-Wallis tests if we could not obtain a normal distribution) to look at the differences of field measured herbivory and herbivore richness between regions and sub-regions. Trait values, fitness and resistance traits, were analyzed using a nested ANOVA (plant family nested within population and population as a factor) to look for differences between the two non-native ranges and the native geographical clades. We further used Tukey post-hoc tests for pairwise comparisons. Lastly, we wanted to test for the predicted tradeoffs between herbivore resistance traits and competitive ability traits in the non-native populations. To narrow down important traits as well as suits of traits we used PCA to find the two most important contributors to variation (components) for resistance traits and then for fitness/competitive ability traits for the two introduced regions. We took these components and used a linear regression (with population means of the components to account for population structure) to look for the relationship between the PCA components for resistance traits and the fitness/competitive ability PCA components. In addition to using the PCA components, we used correlation matrices to look at all pairwise trait tradeoffs (using population means) for each region. All ordinations and MRPPs were run in PC ORD v. 6 (McCune & Mefford, 2016) with all other analysis conducted in R (ver. 3.1.1; R Core team 2013).

## Results

### Herbivores and herbivore communities

We found no evidence of specialist herbivores of *M. guttatus* in the non-native populations of the UK or in ENA. Within both non-native regions, all herbivores found have not been reported to consume plants from Scrophulariaceae *sensu lato*. However, the pool of potential specialist species is greater in eastern North America than in the UK. For example, at several of the ENA sites we observed adults of the specialists *Euphydryas phaeton* and *Junonia coenia* in the proximity of the *M. guttatus* populations, although no caterpillars of these species were found feeding on *M. guttatus* in ENA. Both species feed on plants related to *M. guttatus* and those that share similar phytochemistry (i.e., PPGs) making it possible that they could select these plants for oviposition with their offspring consuming the plants. In contrast we did not find any similar occurrence in the UK populations.

The percent of damaged plants differed between the regions and sub-regions (H= 8.89, DF = 2, p = 0.012, Figure 2A). We found three times fewer plants damaged in the UK than in the native WNA region (Dunns non-parametric comparison p <0.001), while the ENA populations had equivalent levels of field damage to the native WNA region (Dunns non-parametric comparison p = 0.154). The comparisons between the introduced populations and individual native sub-regions showed that the UK populations had significantly less field herbivory than all the native sub-regions except the northern sub-region (Dunns non-parametric comparison p < 0.001 for all except northern), while the ENA populations had equivalent herbivore damage (Dunns non-parametric comparison p < 0.087) compared to all the native sub-regions (H = 18.21, DF = 5, p = 0.003). Herbivore richness did not differ significantly between the UK, ENA, and WNA (F_2.37_= 0.83, p = 0.444, Figure 2B). This was also true when comparing the non-native regions to the native sub-regions (F_5,34_=2.33, p = 0.063).

**Figure 2.**
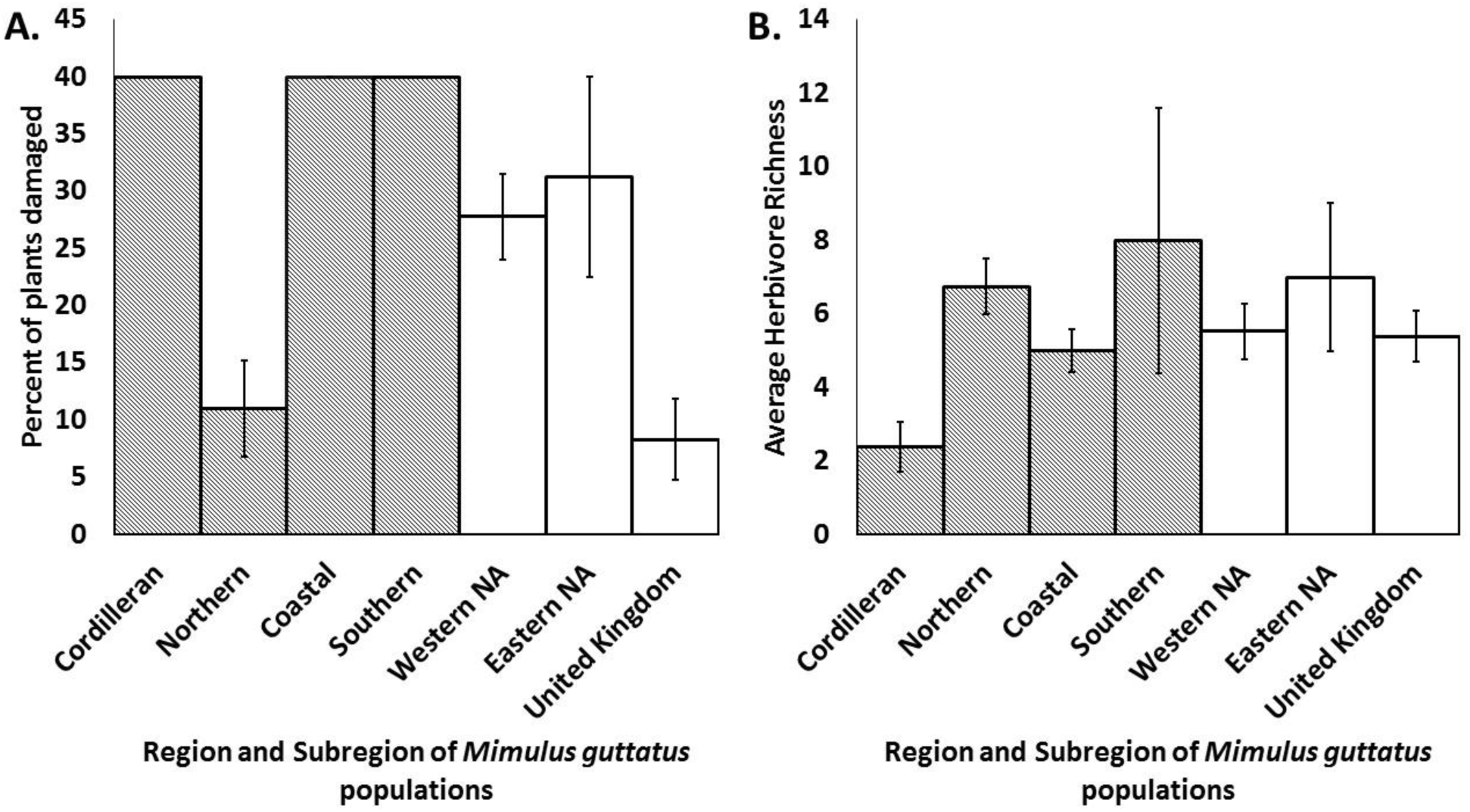
A. Percent of plants with herbivore damage in wild growing *Mimulus guttatus* populations between regions (in white) and native subregions (patterned). B. Average herbivore species richness found in the field feeding on *Mimulus guttatus* populations between regions (in white) and native subregions (patterned). Error bars represent ± 1 standard error. Non-transformed data displayed.

Herbivore communities, at the family level, differed between the native subregions and the non-native populations (MRPP A = 0.085, p < 0.001, Table 1) with the two non-native regions (ENA and the UK) having similar herbivore families to one another (A = −0.017, p = 0.86; Figure 3). The similarity in herbivore communities in ENA and the UK was generally driven by families dominated by generalist herbivores such as terrestrial gastropods and mammals. Differences between the UK populations and the native Cordilleran populations (which includes Alaska and is thus from which the UK populations are thought to be derived; A = 0.092, p <0.001), were driven in part by the lack of leaf mining Agromyzidae in the UK. We also found substantial geographic variation in herbivore community composition within the native subregions. Native subregions were generally separated because of specialist insects that dominated in particular subregions. For instance, leaf mining Agromyzidae flies were common in the Cordilleran subregion as a dominant herbivore while the more southern subregions were dominated by specialist caterpillar species. Herbivore functional feeding guild differences across regions were similar to these herbivore community patterns (Table 1), and were driven by generalist chewers being more common in the non-native regions.

**Figure 3.**
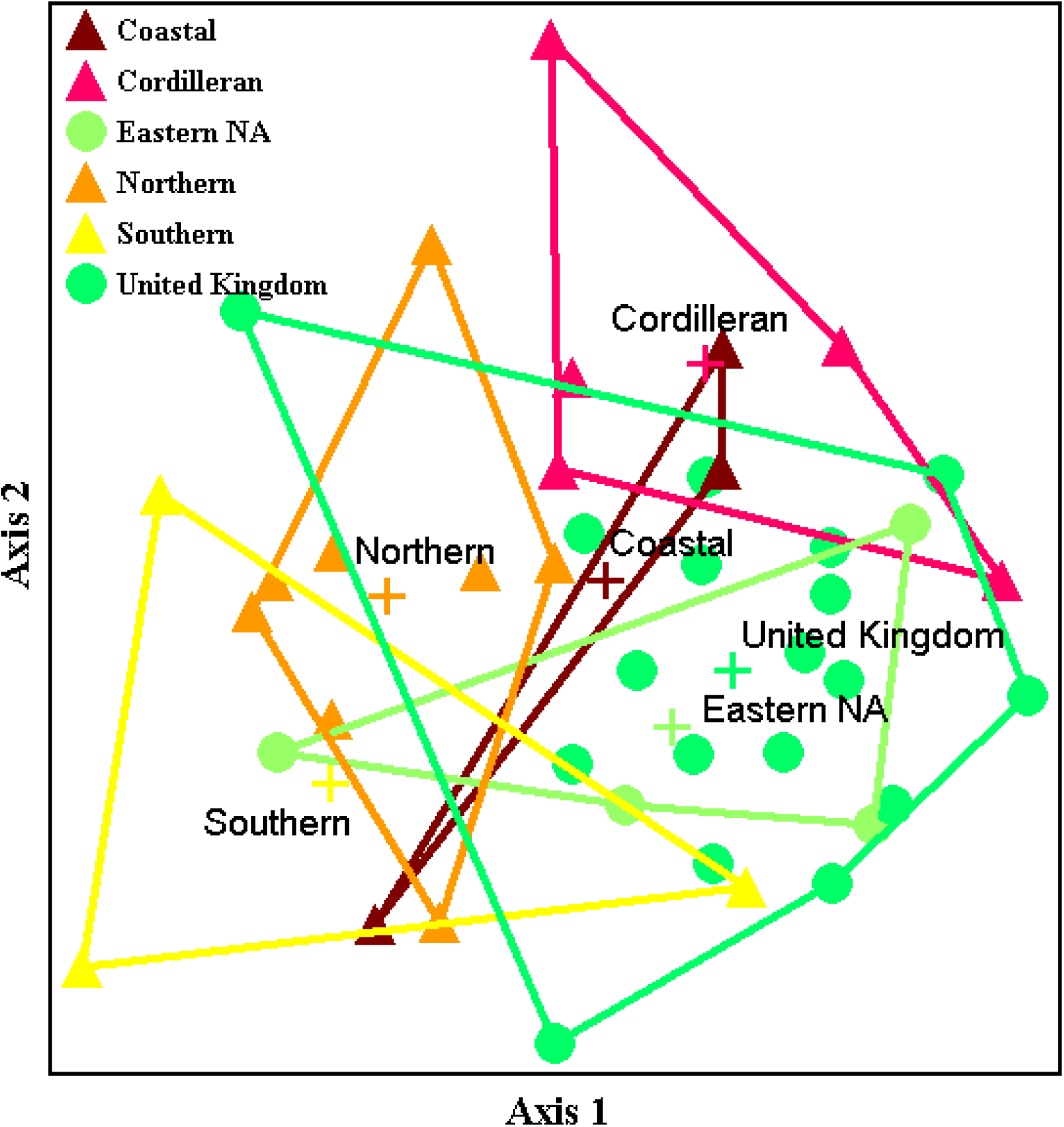
NMDS of herbivore communities based on family for the two non-native regions and the four native subregions. A 2D solution was the best solution with final stress being 25.71. Mean stress of axis 1 was 51.74 and axis 2 was 27.891.

**Table 1.** MRPP results for differences between the non-native regions and the native sub-regions herbivore community at the family level (on bottom and in grey) and functional feeding group (on top in white). The full model was significant for herbivore communities at the family level (A = 0.085, p < 0.001) and for functional feeding groups (A = 0.131, p < 0.001). Bolded results are significantly different pair wise comparisons.

|  | Coastal | Cordilleran | Eastern NA | Northern | Southern | United Kingdom |
| --- | --- | --- | --- | --- | --- | --- |
| Coastal | | $A = 0.083$<br>$p = 0.082$ | $A = -0.063$<br>$p = 0.817$ | $A = -0.062$<br>$p = 0.894$ | $A = -0.069$<br>$p = 0.081$ | $A = -0.029$<br>$p = 0.884$ |
| Cordilleran | $A = 0.020$<br>$p = 0.332$ | | **$A = 0.154$**<br>**$p = 0.019$** | **$A = 0.292$**<br>**$p < 0.001$** | **$A = 0.177$**<br>**$p = 0.008$** | **$A = 0.131$**<br>**$p < 0.001$** |
| Eastern NA | $A = -0.078$<br>$p = 0.908$ | **$A = 0.128$**<br>**$p = 0.0149$** | | $A = 0.066$<br>$p = 0.066$ | $A = -0.031$<br>$p = 0.625$ | $A = 0.014$<br>$p = 0.228$ |
| Northern | $A = -0.042$<br>$p = 0.890$ | **$A = 0.181$**<br>**$p < 0.001$** | **$A = 0.067$**<br>**$p = 0.028$** | | $A = -0.026$<br>$p = 0.680$ | **$A = 0.126$**<br>**$p < 0.001$** |
| Southern | $A = -0.053$<br>$p = 0.812$ | **$A = 0.126$**<br>**$p = 0.019$** | $A = 0.005$<br>$p = 0.433$ | $A = 0.025$<br>$p = 0.215$ | | $A = 0.039$<br>$p = 0.072$ |
| United Kingdom | $A = -0.038$<br>$p = 0.998$ | **$A = 0.092$**<br>**$p < 0.001$** | $A = -0.017$<br>$p = 0.866$ | **$A = 0.099$**<br>**$p < 0.001$** | $A = 0.028$<br>$p = 0.062$ | |

### Herbivore resistance traits

In comparing traits between non-native and native regions, we focus on trait comparisons between populations from the non-native ENA and the native WNA regions and between the non-native UK populations and their likely ancestral WNA Cordilleran subregion.

We found mixed evidence of an overall relaxation of selection on resistance traits predicted by EICA in the non-native *M. guttatus* populations. Physical resistance traits varied between native and non-native regions. Trichome density was significantly different between all regions (F_2,518_ = 86.63, p < 0.001, Figure 4). In support of EICA, native WNA populations had, on average, three and a half times higher trichome density than the non-native ENA plants, which was similar when using the native sub-regions (F_5,516_ = 56.62, p < 0.001, Figure 4). In contrast to the predictions of EICA, the UK population had one and half times higher trichome density than the native Cordilleran sub-region (Tukey post hoc: p = 0.002). Specific leaf area was not significantly different between any of the native and non-native regions (F_2,518_= 1.82, p = 0.121, Figure 4). Leaf water content in the UK populations was slightly higher than the Cordilleran populations and the non-native ENA populations was slightly higher in than the native WNA populations (F_2,517_= 4.53, p = 0.011, Figure 4), suggesting a relaxation in herbivore defense. Leaf dry matter content did not differ significantly across any of the native and non-native regions (F_2.517_ = 0.93, p = 0.392, Figure 4).

**Figure 4.**
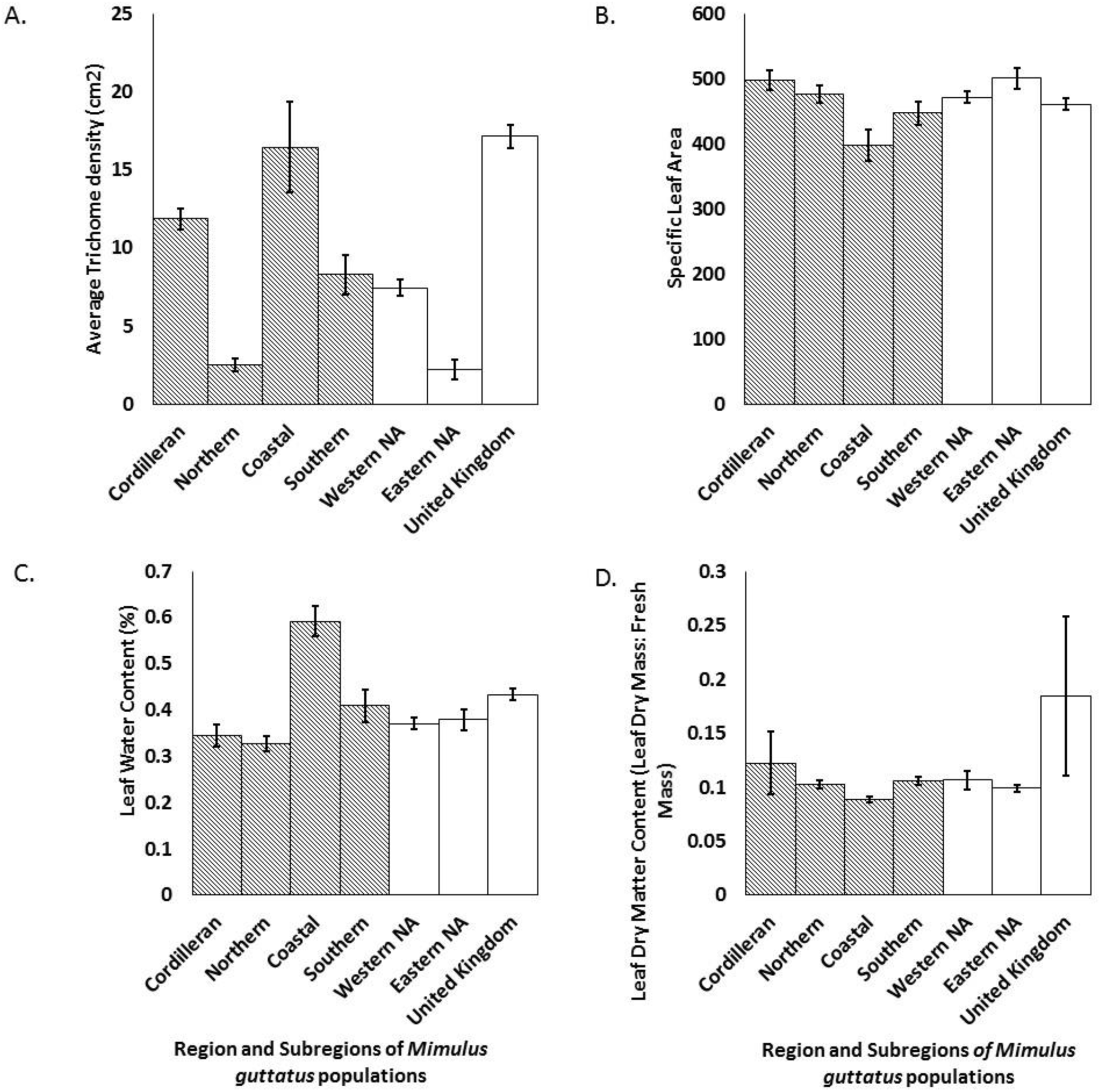
Average physical resistance traits (A. Trichomes, B. Specific Leaf Area, C. Water Content, D. Dry Leaf Matter) in *Mimulus guttatus* populations between regions (in white) and native subregions (patterned). Error bars represent ± 1 standard error. Non-transformed data displayed.

Concentrations of chemical resistance compounds (PPGs) varied across the native and non-native regions (F_2,454_ = 56.62, p = 0.004, Figure 5). Potentially in contrast to EICA, the eastern North American populations had higher levels of total PPGs than the native WNA plants (Tukey post hoc: p=0.004). However, in line with the predictions of EICA the non-native UK plants had lower amounts of total PPG concentration than the native Cordilleran subregion. When considering individual PPGs, there was no consistent overall pattern (Figure 5).

**Figure 5.**
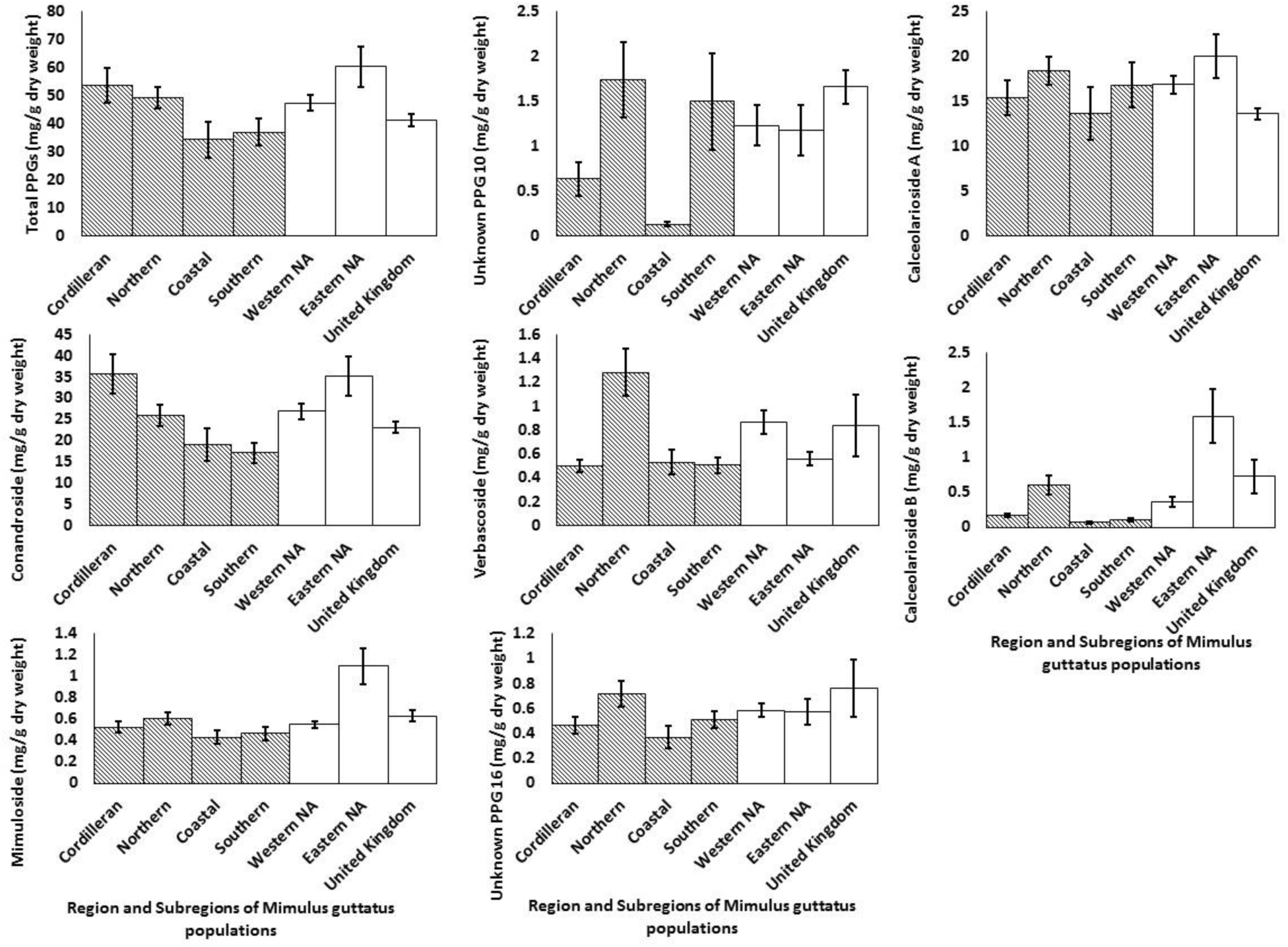
Average concentration (mg/ dry weight) of foliar phenylpropanoid glycosides within regions (white) and subregions (patterned) of *M. guttatus* populations. Error bars represent one standard error. Error bars represent ± 1 standard error. Non-transformed data displayed.

We found no evidence that specialist herbivores performed better on plants from non-native regions than from native, as predicted by EICA. We found no difference in performance of a generalist or a specialist herbivore feeding on tissue from native vs. non-native regions. The generalist caterpillar *Trichoplusia ni* performed similarly on tissue from all regions (F_2,115_ = 0.06, p = 0.940, Figure 6), as well as between the non-native regions and native subregions (F_5,112_ = 1.73, p = 0.131). Performance of the specialist caterpillar *Junonia coenia* also did not differ significantly across native and non-native regions (F_2.41_ = 1.87, p = 0.168). However, there were differences in *J. coenia* performance between the non-native regions and the native subregions (F_5,38_ = 2.77, p = 0.032). Both of the caterpillar species performed equally well on the native Cordilleran subregion plants and the non-native UK plants. Interestingly, the generalist herbivore performed worst on the WNA subregion in which the specialist herbivore species had the highest performance.

**Figure 6.**
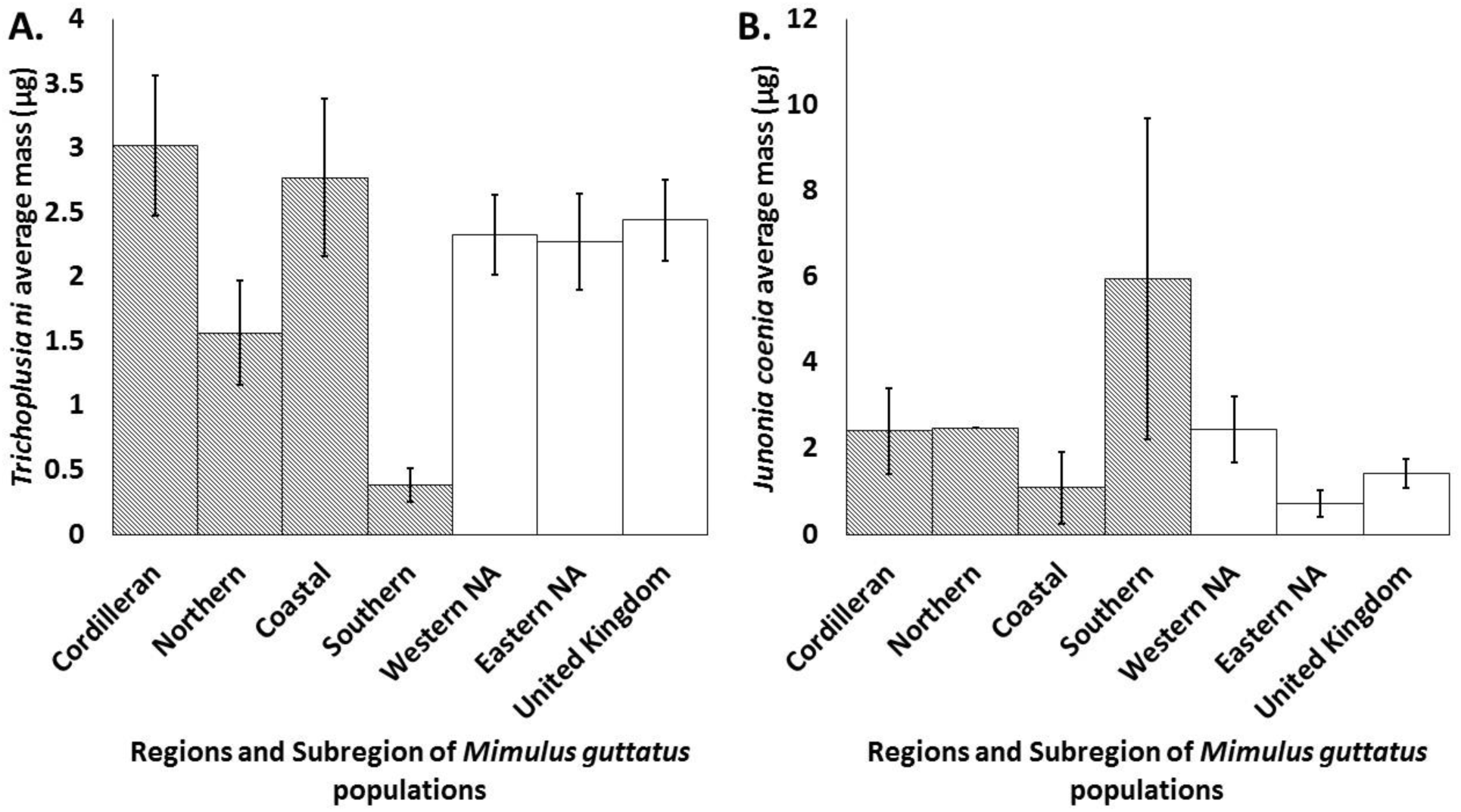
Average performance (mass µg) of (A.) the generalist caterpillar *Trichoplusia ni* and (B.) the specialist caterpillar *Junonia coenia* within regions (white) and subregions (patterned) of *M. guttatus* populations. Error bars represent ± 1 standard error. Non-transformed data displayed.

### Fitness/competitive ability traits

Reproductive traits varied across plants from the native and non-native regions. The non-native ENA populations tended to have relatively equivalent trait values for most traits when compared to the native WNA populations. In contrast, the non-native UK populations deviated from the Cordilleran subregion of WNA for many, but not all traits (Figure 7).

**Figure 7.**
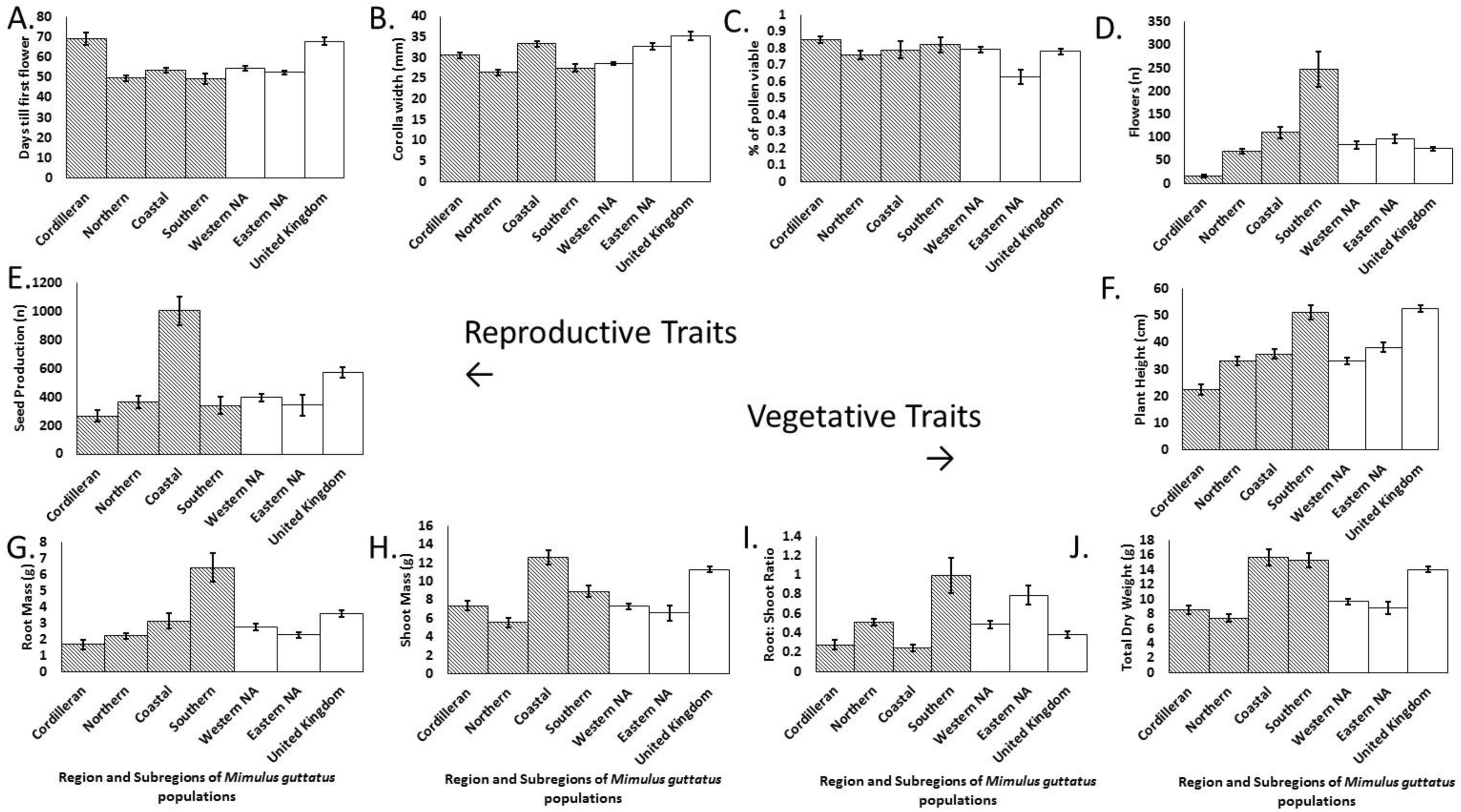
Averages of measures of fitness / competitive ability traits. Reproductive traits: A. Number of days till first flower, B. Width of first corolla, C. Percent of pollen that is viable, D. Total number of flowers produced, E. Number of seeds from first three flowers. Vegetative traits: F. Plant height, G. Root dry mass, H. Shoot dry mass, I. Root:shoot ratio, J. Total dry biomass for *Mimulus guttatus* populations between regions (in white) and native subregions (patterned). Error bars represent ± 1 standard error. Non-transformed data displayed.

Days till flower differed among regions (F_2,481_ = 27.28, p < 0.001, Figure 7). However, the two non-native regions did not significantly differ from their native regions of origin; the ENA plants flowered around the same time as the WNA plants (Tukey post hoc: p = 0.761) and the UK plants flowered at the same time as the Cordilleran subregion plants (Tukey post hoc: p = 0.998). In support of EICA, both non-native regions had on average larger corolla widths than the native WNA region (Tukey post hoc: both p<0.001 compared to WNA plants). This same trend held when comparing the non-native regions to the native subregions (F_5,481_ = 35.83, p <0.001, Figure 7) with the UK plants having larger flowers than Cordilleran plants (Tukey post hoc: p<0.001). While pollen viability was variable across regions (F_2,381_= 8.38, p = 0.003, Figure 7), trends between regions were opposite those predicted by EICA. Pollen viability was lower in the ENA populations than in the WNA as well as in the UK plants versus the Cordilleran plants (need stats for both of these comparisons). Total flower production was variable across regions (F_2,518_= 6.41, p=0.001, Figure 7). Conforming to EICA predictions, the ENA plants produced slightly more flowers on average than WNA plants (Tukey post hoc: p = 0.007), and the UK populations produced on average one and half times more flowers than the native Cordilleran subregion (Tukey post hoc: p>0.001). Finally, seed production varied across regions (F_2,518_= 5.83, p = 0.008, Figure 7). The ENA plants produced an equivalent amount of seeds to the WNA plants (Tukey post hoc: p = 0.064). The UK populations produced twice as many seeds on average compared to the Cordilleran subregion (Tukey post hoc: p <0.001).

For vegetative traits, plants from ENA tended to not conform to the predictions of EICA, while the non-native UK populations did, for most but not all traits. Plant height varied across regions (F_2,502_ = 50.92, p <0.001, Figure 7). Patterns in both non-native regions were compatible with the predictions of EICA. Plants from both the non-native ENA population and the UK population were larger than their native counterparts (Tukey post hoc: p < 0.001), with the UK plants being over twice as tall on average than the native Cordilleran subregion.

There was variation across the regions for total plant biomass (aboveground + belowground; F_2,503_= 47.92, p <0.001, Figure 7), as well as aboveground biomass and belowground biomass considered independently (F_2,497_= 55.36, p<0.001; F_2,501_ = 12.03, p<0.001, respectively). The non-native ENA populations had equivalent total, aboveground, and belowground biomass to the native WNA populations (Tukey post hoc: p = 0.899; p = 0.924; p = 0.941, respectively). As predicted by EICA, the UK populations had almost twice as much total biomass and aboveground biomass, and also higher root biomass than the Cordilleran subregion (Tukey post hoc: p < 0.001 for all biomass comparisons).

Shoot to root ratios varied across regions (F_2,497_= 14.36, p<0.001 Figure 7), with the non-native ENA populations having the largest ratio, which was significantly larger than that for WNA (Tukey post hoc: p = 0.002). Shoot:root ratios for the UK populations were equivalent to those of the Cordilleran populations (Tukey post hoc: p = 0.625).

### Tradeoffs between herbivore resistance traits and fitness/competitive ability

We found some evidence of resistance-fitness/competitive ability trade-offs in the non-native UK region. For herbivore resistance traits, PCA one (24.8%) was associated with chemical traits such as conandroside, calceolarioside A & B, and unknown PPG 16. The second component (15.8%) was associated with trichome density, SLA, and unknown PPG 10. For fitness/competitive traits, PCA one (28.5%) was associated with plant height, number of flowers, and pollen viability while component two was associated with days till first flower, corolla width, and root mass. We found negative associations (suspected tradeoffs) between fitness/competitive ability PCA component one and resistance traits PCA component two (R^2^ = −0.29, p = 0.012; Figure 8). Additionally, we found a positive relationship between fitness/competitive ability component two and resistance component one (R^2^ = 0.17, p = 0.045). The other regressions had non-significant relationships (Table 2).

**Figure 8.**
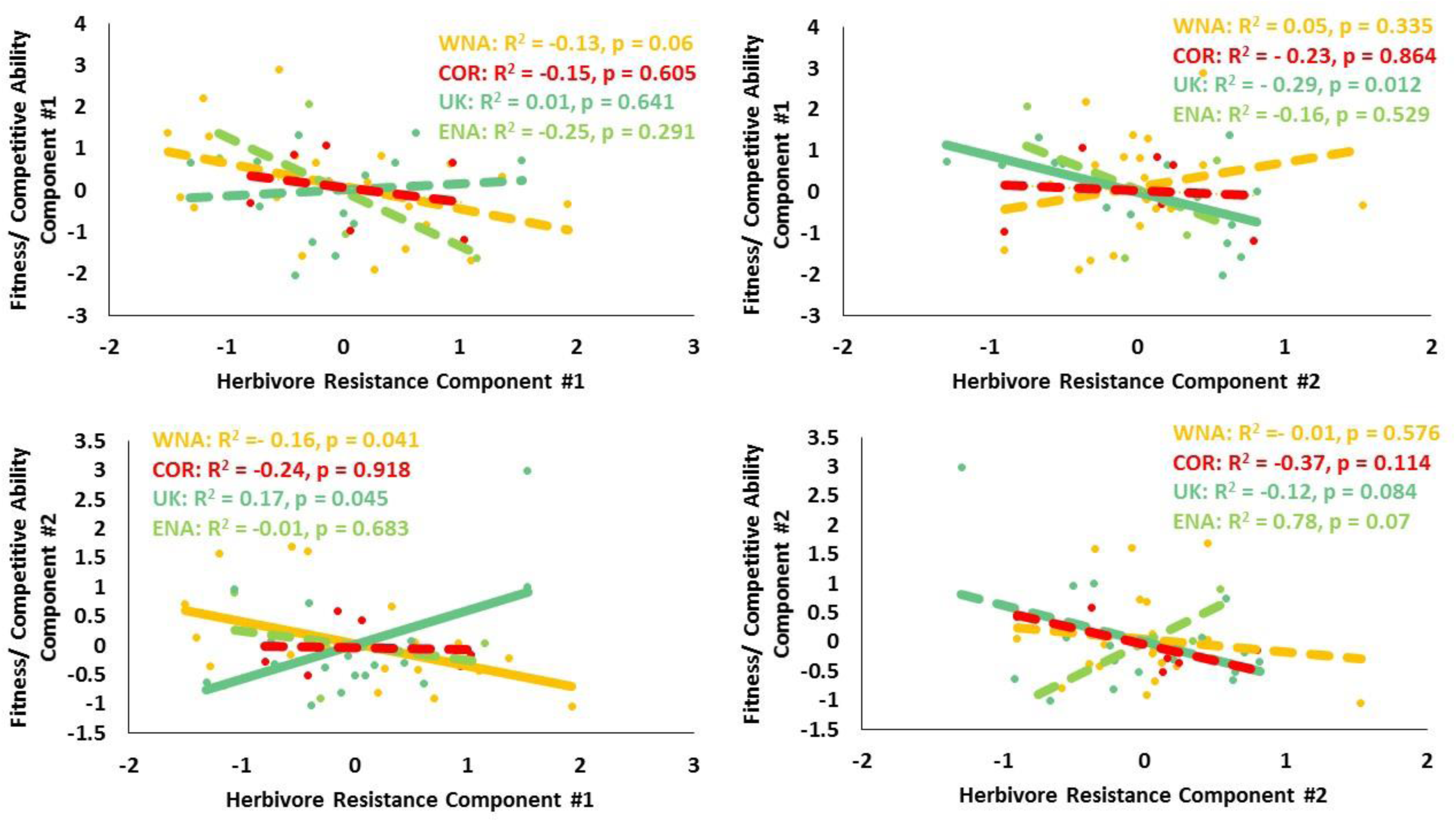
Regressions between fitness/ competitive traits PCA components and herbivore resistance trait PCA components of population means. WNA plants in Orange, Cordilleran plants (COR) in red, UK plants in teal and Eastern North American plants in green. Significant trend lines shown as solid with insignificant trend lines dotted.

**Table 2.** Regression tradeoff results of fitness/ competitive ability PCA components vs herbivore resistance traits PCA components of population means. Significant results are in bold. PCA components are different for each of the regions, and are listed in the text.

|  | Herbivore Resistance Component #1 | Herbivore Resistance Component #2 |
| --- | --- | --- |
| Fitness/ Competitive Ability Component #1 | WNA: $R^2 = -0.13$, $p = 0.06$, $\beta = -0.42$<br>COR: $R^2 = -0.15$, $p = 0.605$, $\beta = -0.26$<br>UK: $R^2 = 0.01$, $p = 0.641$, $\beta = 0.11$<br>ENA: $R^2 = -0.25$, $p = 0.291$, $\beta = -0.71$ | WNA: $R^2 = 0.05$, $p = 0.335$, $\beta = 0.22$<br>COR: $R^2 = -0.23$, $p = 0.864$, $\beta = -0.09$<br>**UK: $R^2 = -0.29$, $p = 0.012$, $\beta = -0.57$**<br>ENA: $R^2 = -0.16$, $p = 0.529$, $\beta = -0.47$ |
| Fitness/ Competitive Ability Component #2 | **WNA: $R^2 = -0.16$, $p = 0.041$, $\beta = -0.45$**<br>COR: $R^2 = -0.24$, $p = 0.918$, $\beta = -0.05$<br>**UK: $R^2 = 0.17$, $p = 0.045$, $\beta = -0.57$**<br>ENA: $R^2 = -0.01$, $p = 0.683$, $\beta = -0.47$ | WNA: $R^2 = -0.01$, $p = 0.576$, $\beta = -0.13$<br>COR: $R^2 = -0.37$, $p = 0.114$, $\beta = -0.71$<br>UK: $R^2 = -0.12$, $p = 0.084$, $\beta = -0.42$<br>ENA: $R^2 = 0.78$, $p = 0.07$, $\beta = 0.92$ |

We found little sign of resistance-fitness/competitive ability trade-offs in the ENA plants. The PCA for resistance traits in the ENA plants had the first component (25%) associated with conandroside, calceolarioside A, and unknown PPG 16 with the second component (20.9%) associated primarily with verbascoside, mimuloside, and unknown PPG 10. The fitness/ competitive traits PCA had a first component (40%) associated primarily with number of flowers, shoot mass, and corolla width. The second component (18.8%) was associated with root mass, pollen viability and seed production. All the components had non-significant relationships to one another (Table 2).

The native region (WNA) also showed evidence of resistance vs. fitness/competitive ability tradeoffs. Herbivore resistance PCA first component (21.2%) was associated with unknown PPG 16, calceolarioside B, and conandroside while the second component (14%) was associated with calceolarioside A, unknown PPG 10, and mimuloside. The first component (33.3%) for fitness/ competitive ability traits was composed primarily of corolla width, plant height, and shoot mass. The second component (16.5%) was associated with days till first flower, number of flowers, and pollen viability. The only significant relationship we found for WNA plants was between fitness/competitive ability component two and resistance component one (R^2^ = −0.16, p = 0.041, Figure 8). All other comparisons were non-significant (Table 2).

Finally, Cordilleran plants showed no signs of tradeoffs. The herbivore resistance PCA component one (25.6%) was associated with unknown PPG 16, calceolarioside A, and conandroside and the second component (16.9%) was associated with calceolarioside B, verbascoside, and unknown PPG 10. The first component of the fitness/ competitive ability traits (41.9%) was associated with corolla width, plant height, and number of flowers produces, the second fitness/ competitive ability component (23.4%) was associated with root mass, seed count, and percent pollen viability. We found no evidence of tradeoffs between these components (Figure 8, Table 2).

## Discussion

By comparing two different plant invasions of differing ages to their native counterparts we found some, but not comprehensive, support for EICA. Support was strongest in the non-native UK, the older of the two invasions. Both the non-native UK and the ENA plants had different herbivore communities than the native WNA plants. However there was adherence to the EICA prediction of a reduction in herbivore damage as well as clear evidence of specialist herbivore escape in only the UK range. We found relatively minor support for the prediction that there would be a decline of herbivore resistance traits in the non-native plants, with some changes in trait values in the non-native vs. native regions, but no differences in herbivore performance in no-choice trials. The UK plants were larger, taller, and produced more seeds and flowers than their native counterparts, in accordance with EICA predictions, while the non-native ENA plants were generally smaller and had poorer pollen production than the native WNA plants. Lastly the UK plants exhibited some tradeoffs between resistance traits and fitness/ competitive ability while the ENA plants did not, confirming to the predictions that release from specialist herbivores can result in allocational tradeoffs that allow for increases in fitness/ competitive ability.

### Enemy release and resistance traits in the non-native populations

We found some evidence of escape from coevolved specialist herbivores in both of the non-native regions. However, this did not translate to the same pattern of relaxed defenses in the two non-native regions. Each non-native region had several resistance traits present at lower levels than in their native ancestral regions. The non-native ENA populations had lower trichome density and higher leaf water content than did the native WNA populations, while the non-native UK populations had higher leaf water content and lower levels of total PPGs than the native Cordilleran region. However, levels of some defenses were also higher in the non-native regions than the native, and we found no difference in performance of generalist and specialist herbivores feeding on native vs. non-native plants. Within a non-native range, even if they are escaping co-evolved specialist herbivores, introduced plants often encounter generalist herbivores that may prefer to attack these non-native plants (Maron & Vilà, 2001; Parker & Hay, 2005; Liu et al., 2007). One of the few other studies that have compared two invaded regions within the context of EICA (other studies have compared multiple invasions but not in the context of EICA) found that populations of the invasive plant *Senecio jacobaea* in a region with a biological control agent (i.e. re-association with a specialist herbivore) did not conform to EICA predictions as well as an invaded region without this control agent present (Rapo et al., 2010). Both non-native ranges in our study had different herbivores communities attacking them than the native region, although herbivory pressures were not necessarily lessened in the non-native environments. Plants in ENA still suffered equivalent damage to WNA plants while, although they suffered less damage, UK plants still had equivalent herbivore richness (per population) as the native plants did.

Although other studies have generally detected EICA-predicted relaxation of resistance to specialist herbivores in non-native regions in feeding trials (Rotter& Holeski, 2018), these changes in herbivore resistance traits were not detected in herbivore performance trials in our study. An alternative hypotheses, the novel weapons hypotheses (Callaway & Ridenour, 2004; Inderjit et al., 2006), predicts that enemy release and non-native success is the result of unique phytochemical compounds or combinations of compounds in the non-native plants that act as deterrents to herbivores in the non-native range. If non-native plants are less palatable than native, but not toxic, this could explain the lack of differences between caterpillar performance in our no-choice trials. The presence of overlap in resistance traits, as some traits likely deter both generalists and specialists, could also result in the overall maintenance of traits that defend against generalist herbivores. The end result would be the maintenance of certain resistance traits that may deter specialist herbivore despite the absence of specialists in the new habitat. For instance, the PPG conandroside has a negative impact on the performance of the generalist herbivores *Grammia incorrupta* and *Spodoptera exigua* as well as a negative impact on the specialist herbivore *Junonia coenia* (Rotter et al., 2018).

### Changes to competitive ability in non-native plants

The EICA prediction that trait values related to fitness and/or competitive ability will be higher in non-native regions was partially supported by our data. Like resistance traits, we did not see similar patterns in fitness and/or competitive ability traits between the two non-native regions. Fitness/competitive ability traits tended not to conform to the predictions of EICA for the non-native ENA region; these trait values were generally very similar to those for the native WNA region. In contrast, fitness/competitive ability trait values were greater in the non-native UK than the native Cordilleran region for many traits, in accordance to EICA predictions. Several other studies have looked at genetic-based phenotypic differences, particularly in physiological and floral traits, between native and non-native *M. guttatus* (van Kleunen & Fischer, 2008; Murren et al., 2009; Martinez, 2018). These studies found that, relative to native plants, the non-native plants produced more flower-bearing stems (but not more flowers; van Kleunen & Fischer, 2008) and had increased flower sizes (Murren et al., 2009). This latter result is similar to our findings in the UK plants. For competitive traits, relative growth rate was not found to be different between native and non-native *M. guttatus* populations (Martinez, 2018). In the UK, *M. guttatus* has been shown to readily spread through both vegetative and seed propagules during high flow events allowing for successful spread (Truscott et al., 2006), although this study focused on non-native populations and did not include a native population comparison.

### Tradeoffs

We found equivocal support for EICA-predicted trade-offs between defense and fitness/competitive ability. In the native WNA and non-native ENA comparison, there were actually fewer detected trade-offs in the non-native region (0) than in the native (1). The native Cordilleran vs. non-native UK comparison was compatible with EICA, with no trade-offs detected in the native region, and one detected in the non-native UK. EICA’s predictions for the success of non-native plants are based on the assumption of allocational tradeoffs existing between herbivore resistance and traits associated with competitive ability (Bloosey & Notzold, 1995; Orians & Ward, 2008). Here, we did find an increase in trait values for traits associated with fitness/competitive ability in one non-native range (the UK), however, these increases were not overwhelmingly associated with decreases in resistance traits. A recent meta-analysis found that non-native plants may in fact not have to make these trade-offs and instead are able to increase resistance traits and fitness/competitive ability (Rotter & Holeski, 2018). This may present some support for hypotheses predicting that non-native plants are able to exploit resources more efficiently or take advantage of unoccupied niche space (Burke & Grime 1996; Davis et al., 2001). In fact there may be a synergy between enemy release and the use of resources as species that are limited by defending themselves may gain a significant advantage when these resources are in abundance (Blumenthal, 2006).

This lack of clear tradeoffs, as predicted by EICA, has also been found in other reviews focused on EICA (Bossdorf et al., 2005; Felker-Quinn et al., 2013). Both of these studies found overall that non-native plant populations changed in their herbivore resistance traits as well as their fitness/ competitive ability traits but these changes did not reflect EICA predictions of a tradeoff (a direct relationship between an increase in fitness/ competitive ability and a decrease in herbivore resistance traits). These studies proposed that more specific looks at relevant traits was needed in testing EICA predictions. Although it is possible that we missed some of the key traits that are involved in tradeoffs, our study was relatively comprehensive in our trait selection particularly for traits important to the ecology of *M. guttatus*.

### Can EICA predict the success of *M. guttatus* invasions?

Finally, our prediction that the more successful invasion (the UK) would display more evidence of adherence to EICA than the less successful invasion (ENA), was supported. The non-native UK populations showed greater adherence to multiple predictions of EICA than the non-native ENA region. Within the EICA framework, species that have become extremely successful invaders such as *Triadica sebifera* (Huang et al., 2010; Carillo et al., 2014,) might conform more closely to EICA than relatively non-invasive non-natives such as *Lepidium draba* (Cripps et al., 2009). In the UK, *M. guttatus* has successfully spread throughout the country, filling many of the available niches. In ENA the invasion is thought to be more recent, *M. guttatus* has become extirpated from several of the locales where it has previously been reported, and no new populations have been reported since at least the early 2000’s. Our results correspond with those of other studies that compared different non-native plant species within the same region that had differing level of invasiveness (ability to spread and dominate communities). Plants that were ranked as more invasive had lower rates of herbivory than those non-natives that were not considered as invasive (Cappuccino & Carpenter, 2005). This supports the idea that the strongest evidence for EICA may be found in more successful invasions.

There are many different frameworks for understanding the success of non-native organisms (Catford et al., 2009) and it is likely that there is not a single one that can consistently and fully explain why a non-native species becomes successful across systems (Gurevitch et al., 2011; Lau & Schultheis, 2015). This is the case with our results; although we found some evidence to support EICA, particularly in the non-native UK region, there were several patterns that were not necessarily compatible with EICA (e.g., caterpillar performance was not different between the native and non-native plants and the sometimes positive relationship between resistance traits and fitness/ competitive ability in the UK plants). However, we do present evidence that the release from (or at least a shift in herbivore suites) can lead to evolutionary changes in plant resistance traits that result in an increase in competitive ability.

## Acknowledgments

We would like to thank the members of the Holeski lab group for assistance on various parts of this project. Phil Patterson helped with plant cultivation in the greenhouse. Thanks to Dean Robinson who let us study the plants on his property. Funding was provided by the NAU genes to environment fellowship program and the NAU Department of Biological Sciences. A special thanks to funding from the Michigan Botanical Foundation, Idaho Native Plant Society, and the Utah Native Plant Society.

## Authors Contributions

M.C. Rotter designed the study, collected data, did the statistical analysis and wrote the manuscript. M. Vallejo-Marin co-wrote the manuscript, helped in field work, and provided background. L.M. Holeski participated in the design for the study, assisted with sampling techniques, and co-wrote the manuscript.

## Data Accessibility

Data will be archived in Dryad.

## Supplemental Information

**Table S1.**
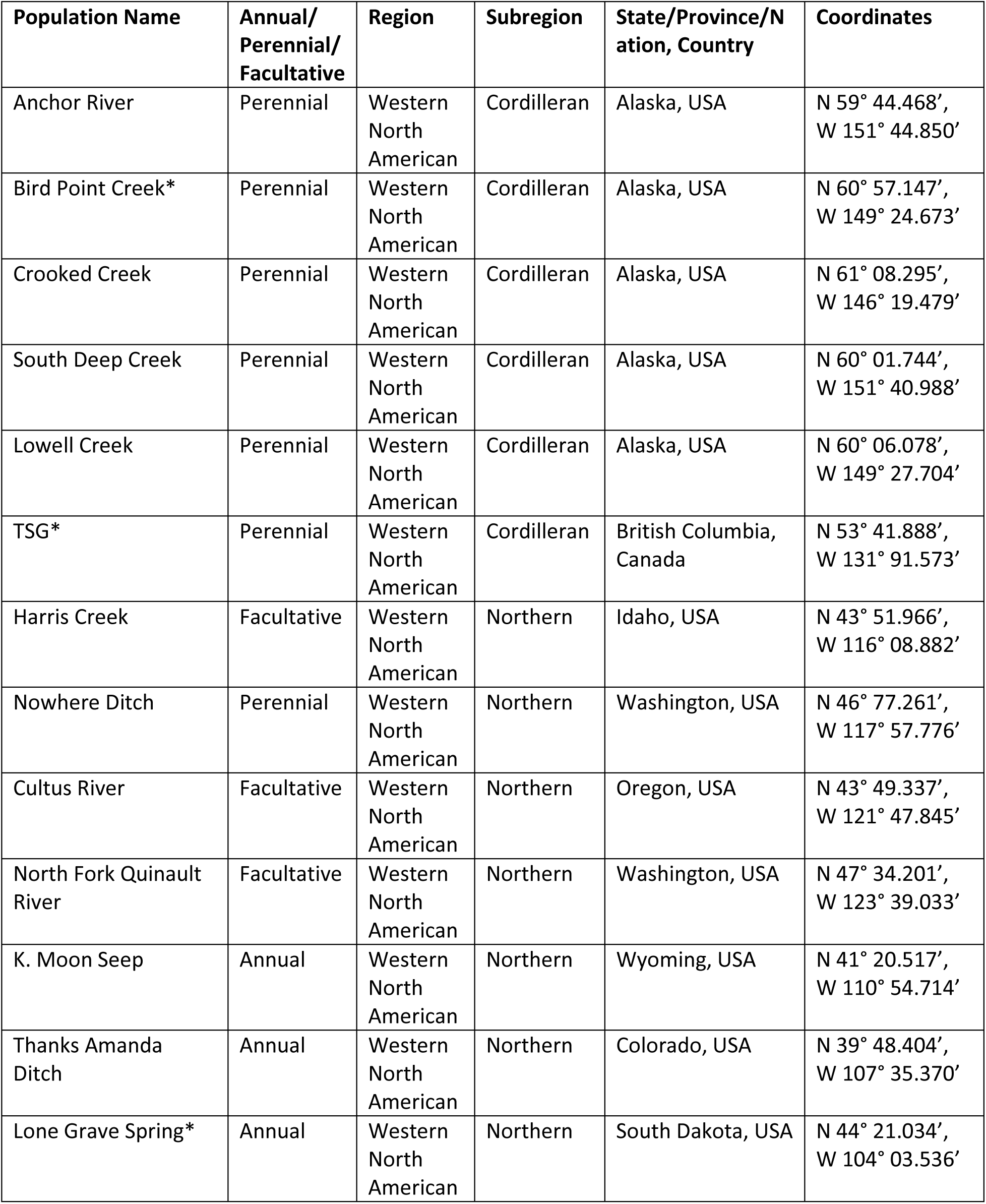
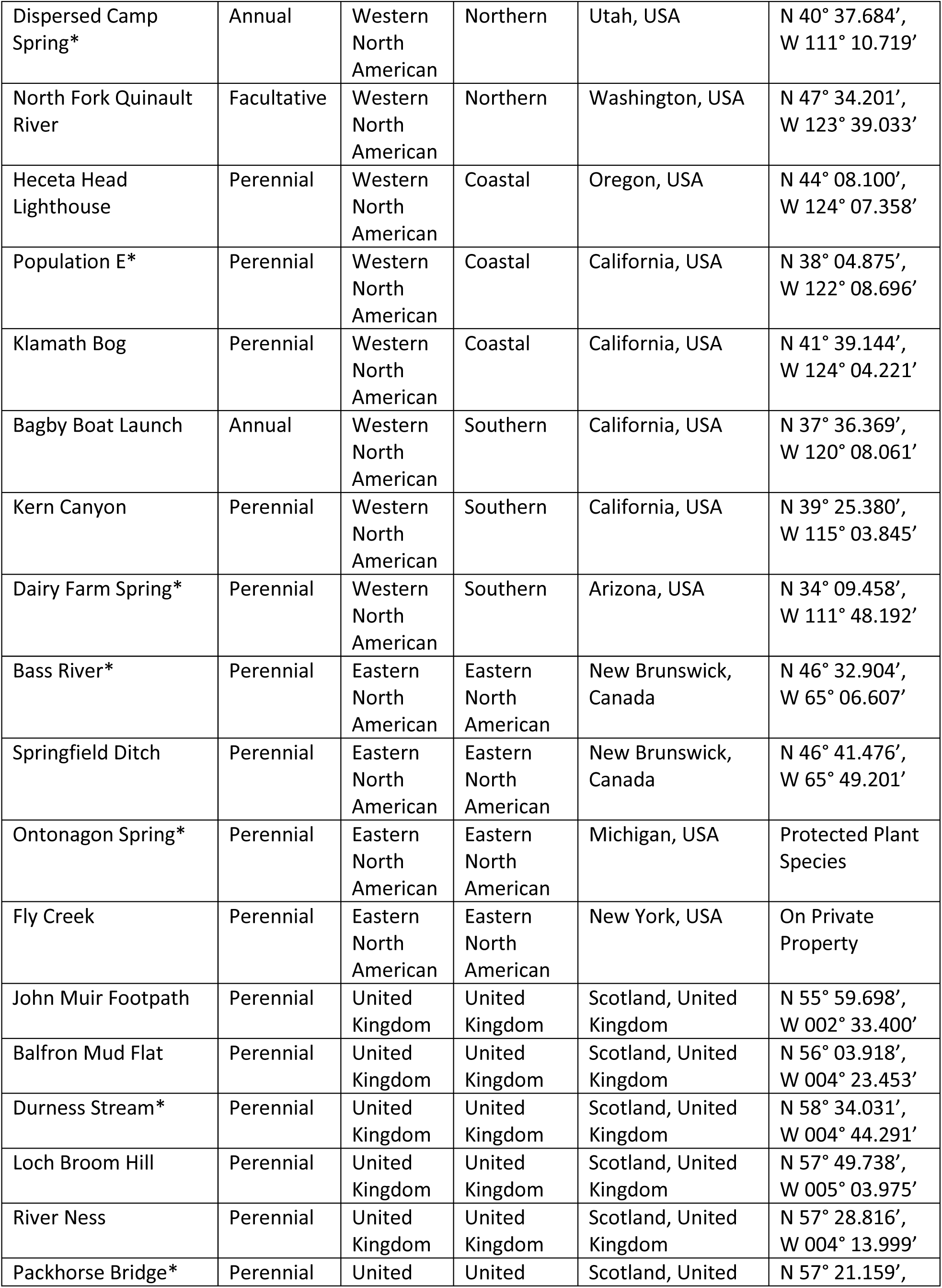
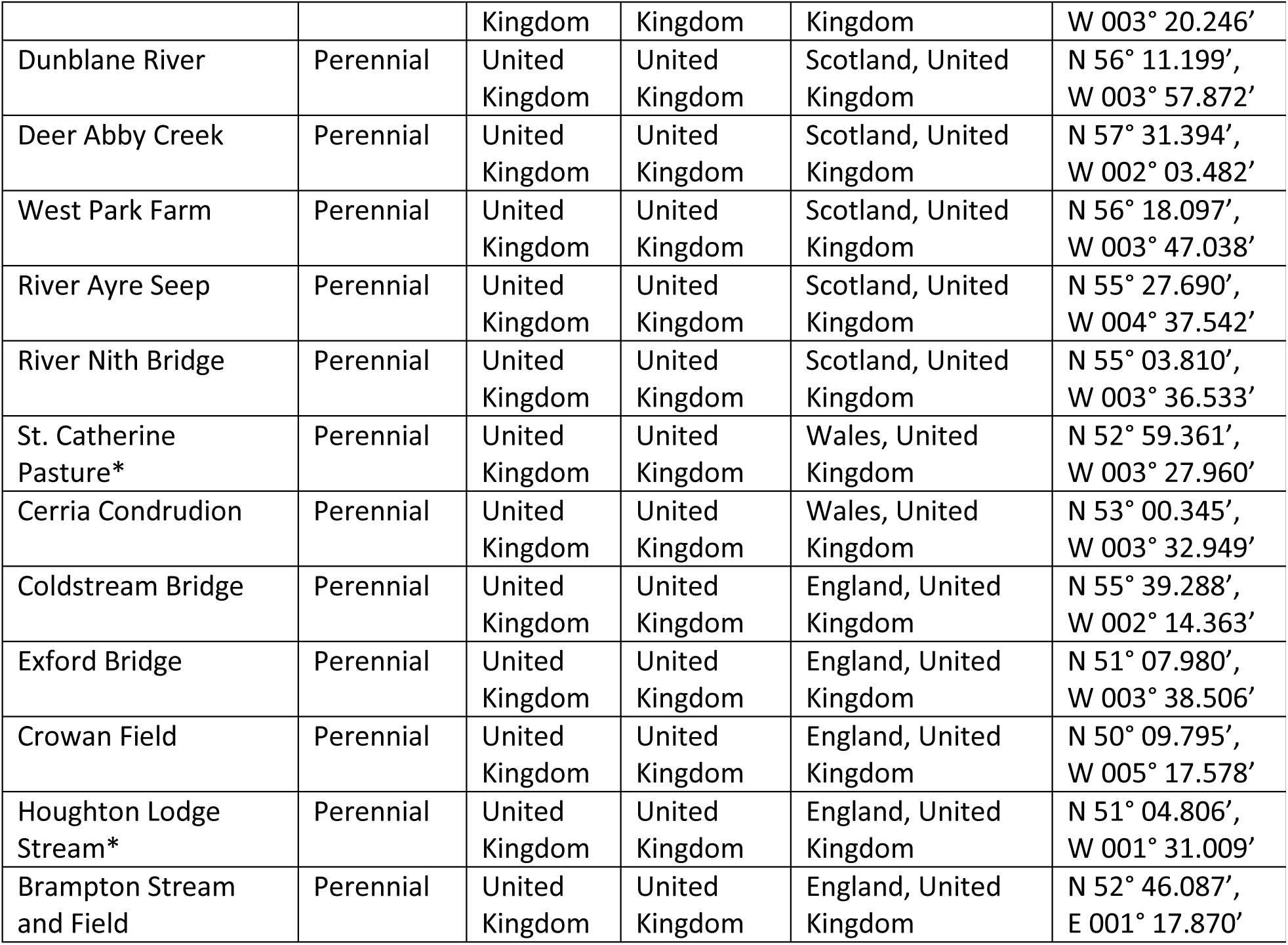
Locations of all populations used in this study. With plant life history, region and subregion (Stace 2010, Twyford and Freidman 2015). Populations with an * were additionally used for caterpillar feeding trials.

**Figure S1.**
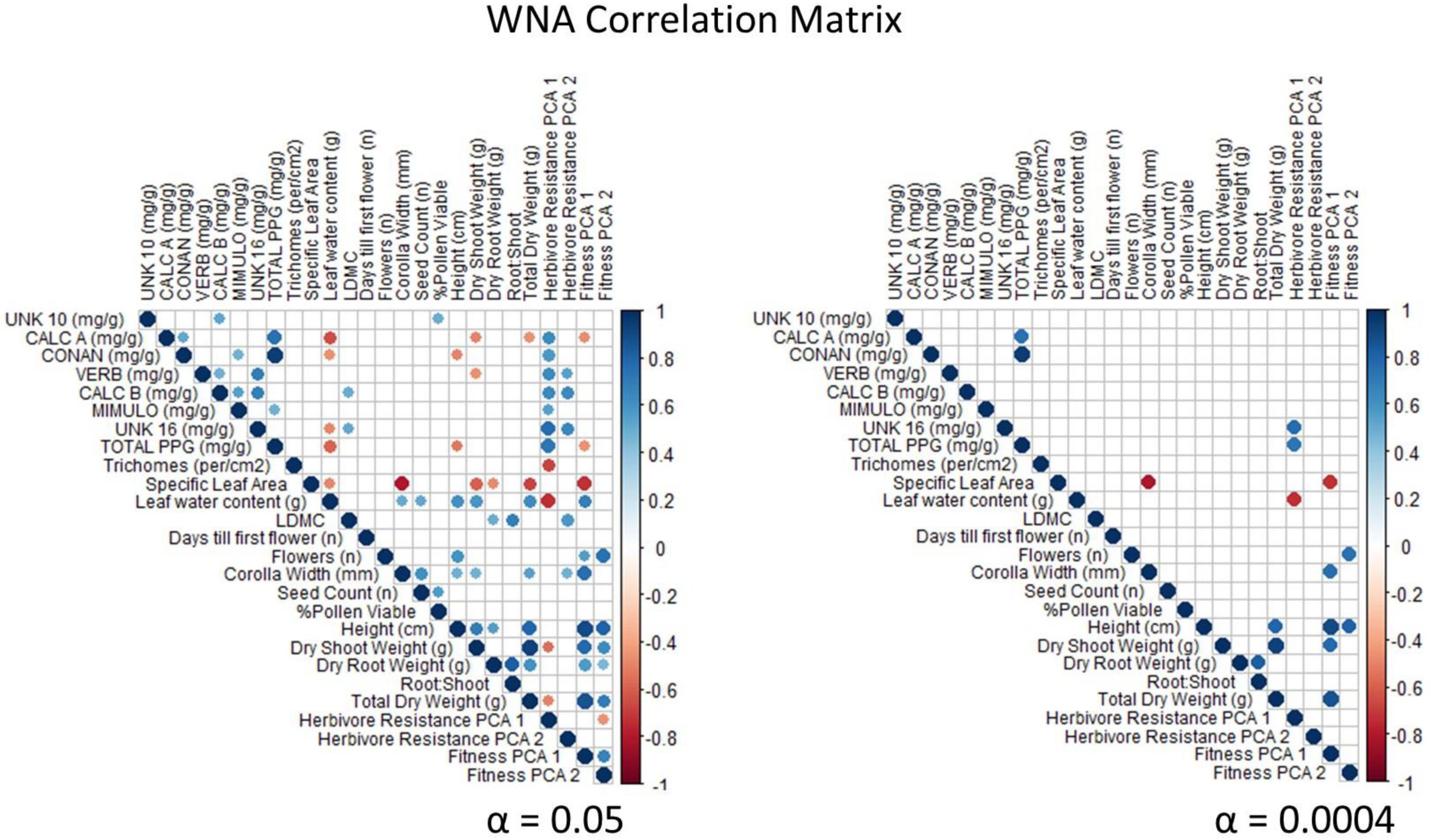
Correlation matrix for population means of all continuous pairwise traits measured for native western North American populations (WNA). Blue indicates a positive r value and red being a negative r value. Only significant r values are displayed. Left figure is with α set at 0.05 and right figure is adjusted α of 0.0004 for multiple tests.

**Figure S2.**
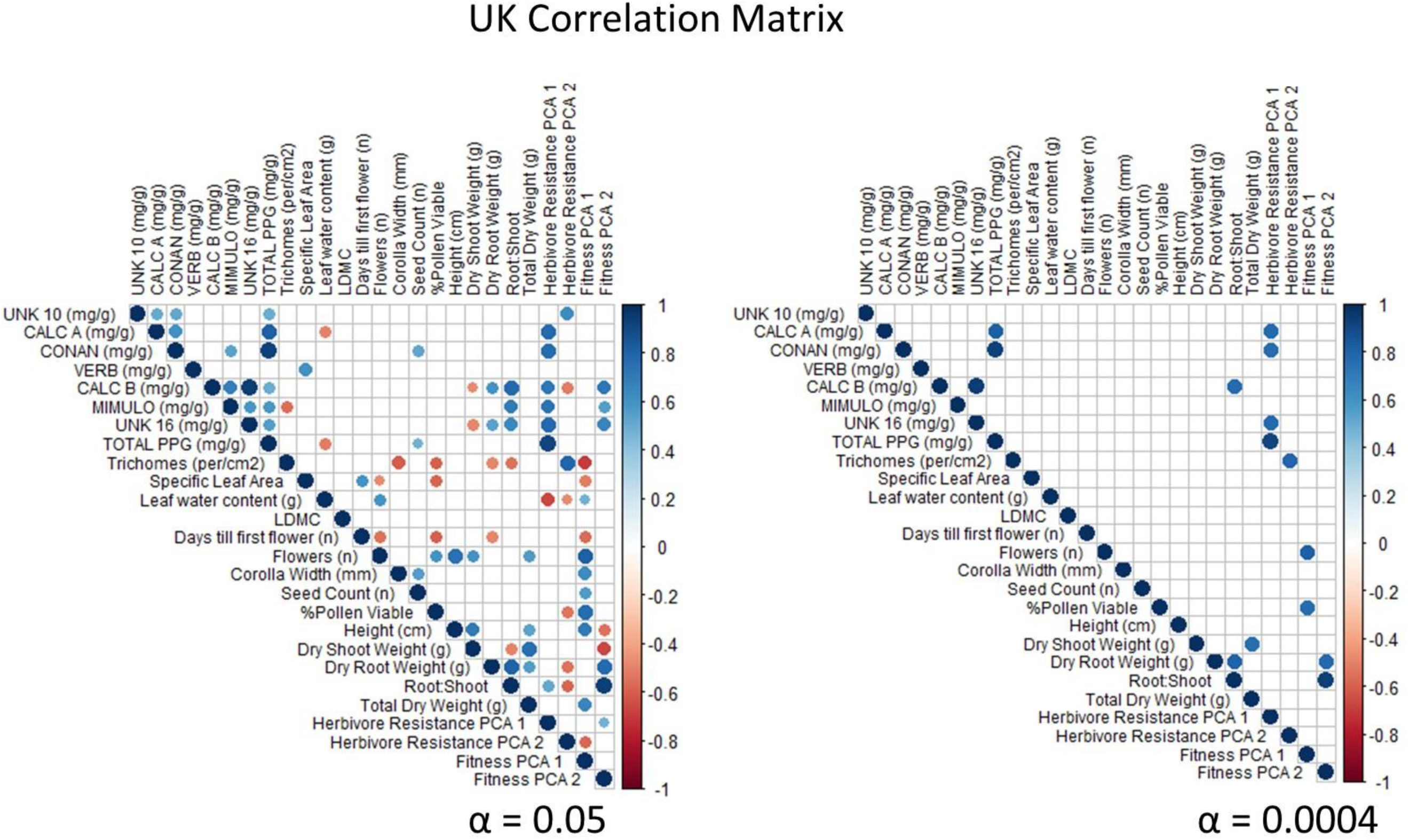
Correlation matrix for population means of all continuous pairwise traits measured for United Kingdom populations (UK). Blue indicates a positive r value and red being a negative r value. Only significant r values are displayed. Left figure is with α set at 0.05 and right figure is adjusted α of 0.0004 for multiple tests.

**Figure S3.**
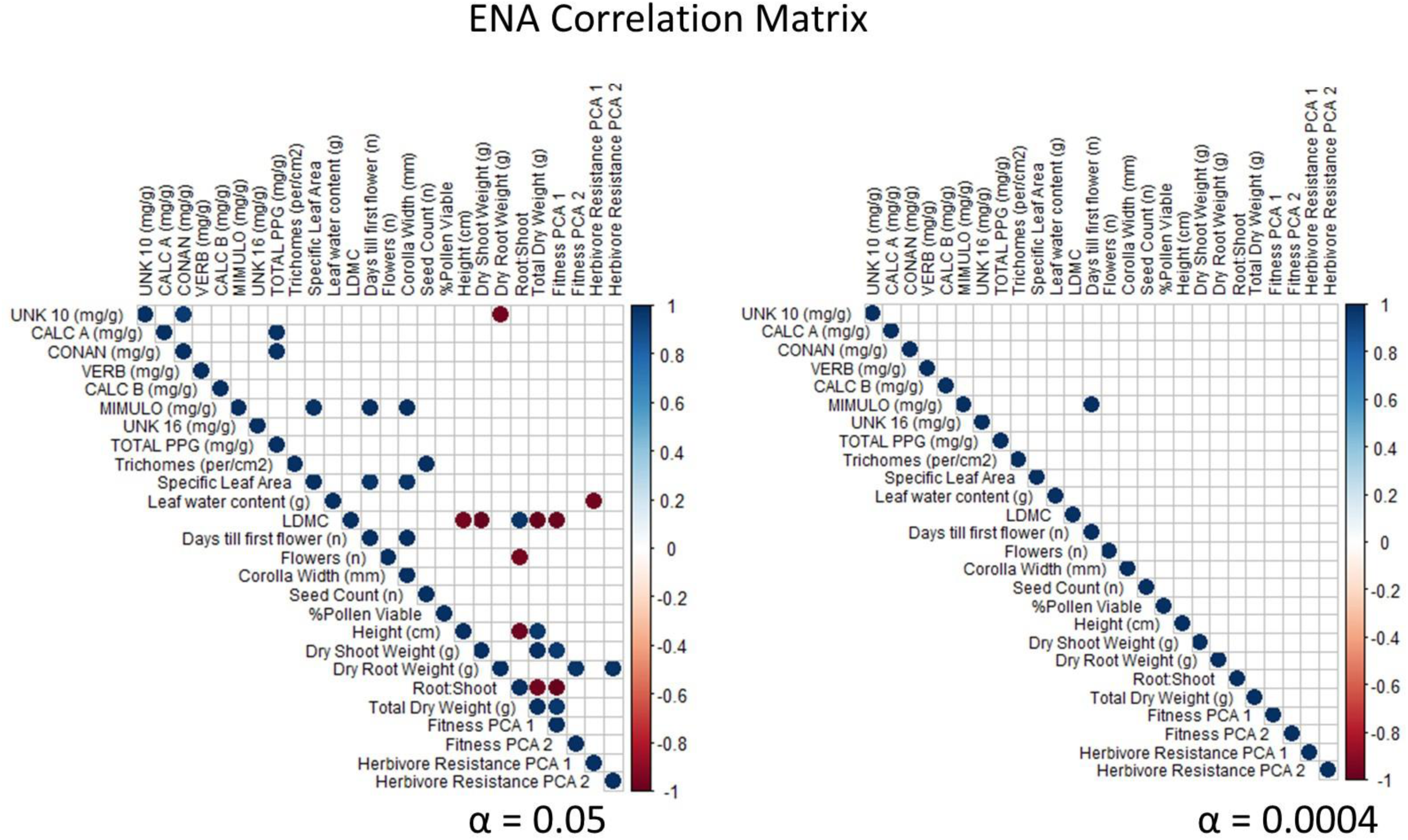
Correlation matrix for population means of all continuous pairwise traits measured for eastern North American populations (ENA). Blue indicates a positive r value and red being a negative r value. Only significant r values are displayed. Left figure is with α set at 0.05 and right figure is adjusted α of 0.0004 for multiple tests.

